# On the potential of Angiosperms353 for population genomics

**DOI:** 10.1101/2020.10.11.335174

**Authors:** Madeline Slimp, Lindsay D. Williams, Haley Hale, Matthew G. Johnson

## Abstract

Targeted sequencing using Angiosperms353 has emerged as a low-cost tool for phylogenetics, with early results spanning scales from all flowering plants to within genera. The use of universal markers at narrower scales—within populations— would eliminate the need for specific marker development while retaining the benefits of full-gene sequences. However, it is unclear whether the Angiosperms353 markers provide sufficient variation within species to calculate demographic parameters. Using herbarium specimens from a 50-year-old floristic survey of Guadalupe Mountains National Park, we sequenced 95 samples from 24 species using Angiosperms353. We adapted a data workflow to process targeted sequencing data that calls variants within each species and prepares data for population genetic analysis. We calculated genetic diversity using standard metrics (e.g. heterozygosity, Tajima’s D). Angiosperms353 gene recovery was associated with genomic library concentration, with limited phylogenetic bias. We identified over 1000 segregating variants with zero missing data within 22 of 24 species. A subset of these variants, which were filtered to remove linked SNPs, revealed high heterozygosity in many species. Tajima’s D calculated within each species indicated a moderate number of markers potentially under selection and identified evidence of population bottlenecks in some species. Despite sequencing few individuals per species, the Angiosperms353 markers contained sufficient variation calculate demographic parameters. Larger sampling within species will allow for estimating gene flow and population dynamics in any angiosperm. Our study will benefit conservation genetics, where Angiosperms353 provides universal repeatable markers, low missing data, and haplotype information.

## INTRODUCTION

### Universal Target Capture

The estimation of demographic parameters in natural populations is a critical tool for species delimitation (Duminil and Di Michele, 2009), biogeography (Overcast et al., 2019), and monitoring of populations and species in a dynamic changing environment (Allendorf et al., 2010). The feasibility of estimating demographic parameters (including heterozygosity, effective population size, and levels of introgression) in non-model taxa relies on markers that can detect sufficient variation across the genome while remaining cost-effective for analysis on hundreds of individuals. In plants, population genomics could benefit from markers that enable the further unlocking of herbarium specimens for botanical research, following advances in phylogenomics (Shee et al., 2020), microbiome research (Heberling and Burke, 2019), and the effects of climate change on plant populations (Miller-Rushing et al., 2009).

Traditional Sanger sequencing of PCR amplicons often employ universal primer sequences, but the genes targeted (e.g. the plastid markers matK & rbcL, or the nuclear ribosomal ITS) do not contain sufficient variation to allow species differentiation, meaning population level analysis is unlikely to provide conclusive results (Supple and Shapiro, 2018). Whole genome methods are also not feasible for many plant species, which can have large genome sizes (e.g. Fritillaria, 86 GBp) (Kelly et al., 2015). Genome skimming, including extraction of organellar genomes and the ribosomal cistron, is popular for phylogenetics of non-model organisms (McKain et al., 2018). However, organellar regions do not represent sufficient numbers of unlinked sites to generate unbiased estimates of genetic diversity (McMahon et al., 2014). More recent efforts have focused on optimization of Reduced Representation Sequencing, including Restriction-digest Associated DNA sequencing (RADseq), RNA sequencing (RNAseq), and target capture via hybrid enrichment (HybSeq). Sequencing RNA of non-model species, while generating data for tens of thousands of genes, is still prohibitively expensive at the population level and its use is limited in conservation genomics. In contrast, RADseq (and variations such as Genotyping by Sequencing, double-digest RAD sequencing, etc.) has gained popularity in population genetics because the method requires no primers and may be used in any organism to potentially generate thousands of markers (Andrews et al., 2016). However, RADseq without a reference genome requires that short sequences must be clustered by sequence similarity across samples to identify loci (Eaton and Overcast, 2020), often generating large amounts of missing data.

In contrast, target capture sequencing of exons and flanking non-coding regions (HybSeq, Weitemier et al., 2014) holds several advantages for estimating demographic parameters. Target capture methods generate data for hundreds of loci at a reasonable per-sample cost (Hale et al., 2020), result in datasets that often have limited missing data (Carter et al., 2019), and are suitable for degraded DNA such as herbarium specimens (Brewer et al., 2019). The development of universal probe sets for HybSeq across large groups (Buddenhagen et al., 2016; Johnson et al., 2018) further suggests a role for HybSeq at the population level, if the loci are variable.

### The GUMO Collection as a Test Case

Located in the Trans-Pecos region in far-west Texas, the Guadalupe Mountains National Park (GUMO) has been described as the “most botanically diverse area of Texas.” (Eason, 2018). The park contains the basin-range system of the Chihuahuan Desert, the world’s most extensive Permian fossil reef, and abrupt elevation changes (including the Guadalupe Peak at 2667 m). For 100 years prior to the National Park opening in 1973, it was used for agriculture, including livestock husbandry. Goats grazed the landscape of the park, heavily disturbing the natural population vegetation balances (Glass et al., 1974). The Guadalupe Mountains are isolated from other high-altitude peaks, which may impact the persistence of many plant species in a warming climate. Characterizing the impact of both land-use and climate change is an important goal for the Park, and ideally would include conservation genomics on species of concern (Allen, 2013). Northington and Burgess (1976) made the first comprehensive floristic survey of the newly created National Park, collecting an estimated 3000 specimens from over 400 species. The Guadalupe Mountains are an interesting botanical region due to geologic variabilities and intersection of three ecoregions: desert scrublands, high plains, and montane forest at higher elevations (Figure 1). Northington and Burgess described 14 species as endemic to a limited area in and/or around GUMO, with an additional 37 species described as having small ranges with their furthest extent in GUMO. Their collection, now housed at the E.L. Reed Herbarium at Texas Tech University (Herbarium Code: TTC), includes 55 species with at least 6 unique specimens, raising the potential for estimating within-species demographic parameters. The extensive collection, not digitized until recently, provides a unique snapshot of a past botanical community, especially for rare species such as *Philadelphus hitchcockianus* and *Salvia summa* (Figure 1).

**Figure 1:**
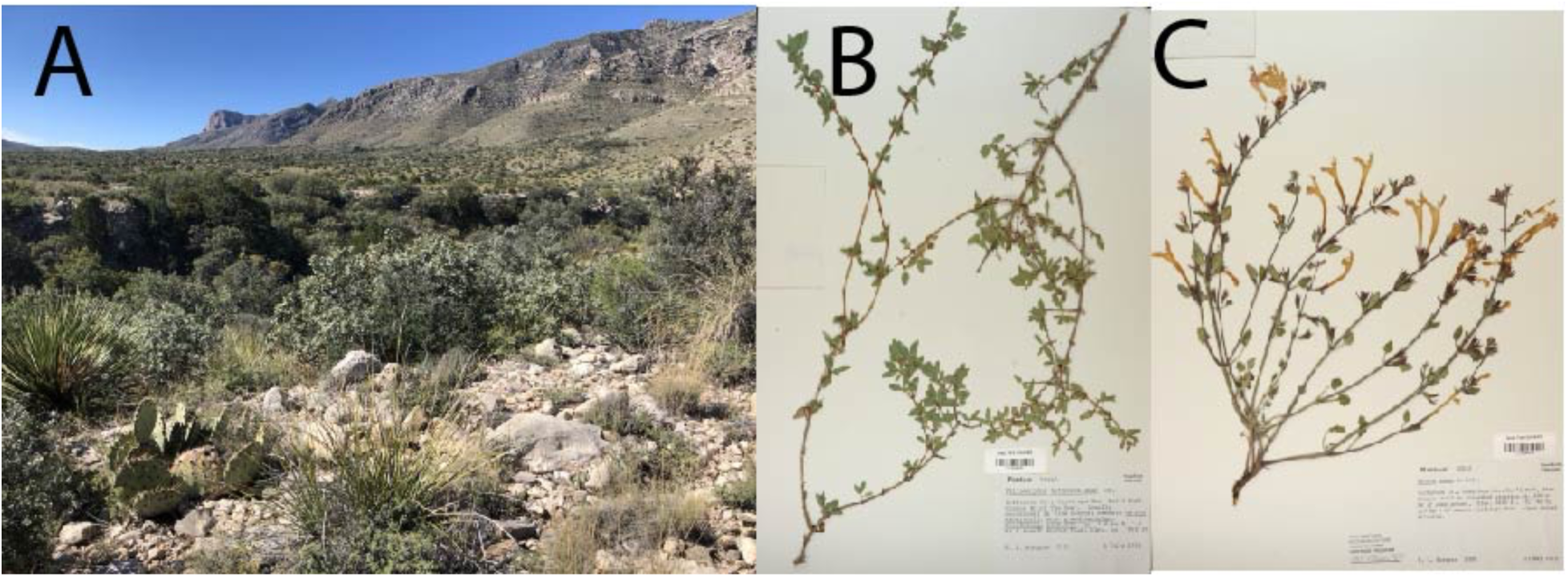
A: Typical desert scrubland habitat in Guadalupe Mountains National Park (GUMO). B and C: Representative herbarium specimens from the 1970s GUMO collection, *Philadelphus hitchockianus* (B) and *Salvia summa* (C), two species with ranges restricted to the GUMO region.

In this study we aim to establish a universal and cost-effective strategy for recovery of Angiosperms353 genes from the GUMO Collection. Determining whether these changes had effects on the genetic diversity of populations requires molecular markers that can work across species and can be used with herbarium specimens. We also aim to determine whether there is sufficient variability amongst Angiosperms353 genes to calculate population genetics parameters. The Angiosperms353 target capture kit is especially suitable for use with the GUMO collection, because it offers high gene recovery with degraded DNA (Brewer et al., 2019) from any angiosperm genus and is a high output solution that produces whole gene sequences (Johnson et al., 2018). Here, we focus on analyzing recovered gene data from the GUMO collection and calculating inbreeding coefficients, heterozygosity, and Tajima’s D to demonstrate whether Angiosperms353 is suitable for larger population genomics projects.

## MATERIALS AND METHODS

### Sampling

We sampled species based on availability of at least four distinct specimens per species, with a goal of sampling approximately 1 cm2 of tissue from 24 species: 8 grasses, 6 forbs, 6 shrubs, and 4 tree species (Appendix 1). We extracted DNA in 1.1 mL round bottom tubes, to which we added tissue and two steel bearings. We froze the tissue in a SPEX CryoStation, then ground the tissue using a SPEX GenoGrinder MiniG tissue homologizer processing all 95 specimens simultaneously. Due to the diverse species representation on the plate, grind quality differed between samples and was poor in those with extremely fibrous or glabrous tissue. We repeated freezing and grinding twice more until a powder was observed for all specimens. We then proceeded with a typical CTAB/Chloroform DNA extraction method (Doyle and Doyle, 1987) with three modifications: 0.4% 2-mercaptoethanol added to the CTAB buffer, incubation in CTAB for 10 hours, and precipitation of DNA in isopropanol in a −20°C freezer for five days. These modifications have been previously shown to maximize the yield of DNA from herbarium specimens (Brewer et al., 2019). Samples where DNA yield exceeded 5 ng/µL (via Qubit fluorometer) with most fragments above 500 bp (via agarose gel) were sheared using the fragmentase enzyme (New England Biosciences) with a desired fragment size of 500 bp. We did not shear extracts with a DNA concentration or size distributions below either threshold, and instead were concentrated the DNA using a SpeedVac centrifuge.

### Enriched Library Preparation

Using NEBNext Ultra II DNA kit (New England Biosciences) we prepared dual-indexed Illumina sequencing libraries with all reagents in half-volume, targeting an insert size of 500-700 bp. Before library preparation, samples were either diluted or concentrated to achieve an input DNA concentration of ~200 ng in 25 uL. Following final PCR (8 cycles) and cleanup of libraries with homebrewed SPRI beads (Rohland and Reich, 2012) we assessed concentration using a Qubit fluorometer. We measured fragment size distribution for libraries with a concentration less than 5 ng/µL using Agilent TapeStation 2200 on fifteen libraries, to reduce the cost of assessing all 95 libraries. The samples chosen for the TapeStation were selected based on (1) concentration and (2) placement of the library on the plate. Target capture of the Angiosperms353 loci occurred in four separate reactions, each of which contained 24 libraries pooled by their final concentration (Pool 1: 0.139-10.8 ng/µl, Pool 2: 10.9-15.1 ng/µl, Pool 3: 15.4-22.9 ng/µl, Pool 4: 23.1-39.3 ng/µl), regardless of species. Library pools were then concentrated to 7 uL by using a SpeedVac (on low heat) until all liquid was out of the tubes and then re-eluting the libraries in nuclease-free water. Target capture reactions followed Arbor Biosciences’ myBaits Hybridization Capture for Targeted NGS manual version 4.01, except for a 3:1 dilution of the RNA probes compared with manufacture recommendations. This modification has proven efficient for pooling 96 samples in previous targeted sequencing studies (Hale et al., 2020). Hybridization with the Angiosperms353 probes took place at 65°C w for 26 hours, followed by PCR enrichment of the approximately 500-bp-insert libraries for 14 cycles. Each pool was analyzed on the TapeStation and the pools were combined in equimolar fraction (10 nM / pool) before sequencing on the Illumina MiSeq v3 2×300 platform at the Texas Tech University Center for Bioinformatics and Genomics.

### Data processing

We first processed sequence reads using Trimmomatic (Bolger et al., 2014) to remove adapter sequences and reads with an average Q score below 25. Angiosperms353 genes were recovered from cleaned Illumina reads using HybPiper (Johnson, Gardner, et al., 2016). Briefly, the workflow identified potential matches between reads and target genes (using BLASTX, Camacho et al., 2009) to allow for high phylogenetic distance between targeted sequences and sample reads), sorted the reads by gene, and conducted de novo assembly separately on each gene, mitigating the need for a reference sequence for non-model organisms. We used a target file containing representative amino acid sequences of the Angiosperms353 loci obtained from github.com/mossmatters/Angiosperms353. Two types of sequences were output from HybPiper: coding sequences corresponding to the targeted genes, and “supercontigs” that contain both the exon sequences and the flanking non-coding (i.e. intron and other untranslated regions) sequence. To test for genetic diversity in each species, we generated a set of reference sequences by selecting the longest supercontig recovered for that species for each gene (Fig. 2).

**Figure 2:**
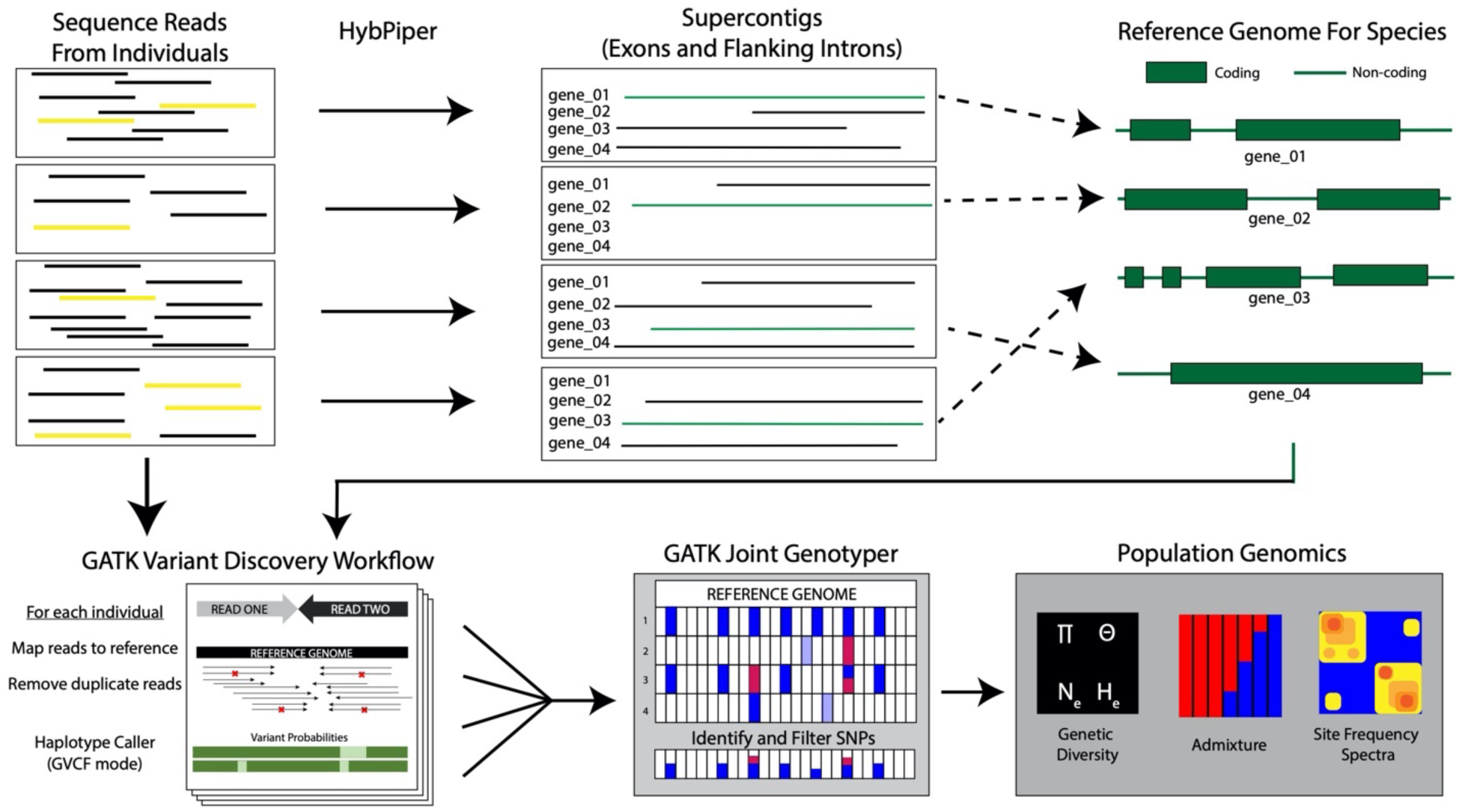
Data Workflow for identifying SNPs within each species from target capture data. Reference genomes for each species were constructed by selecting the longest supercontig from each gene for each species.

### Variant Detection and Filtering

To detect variants within species we followed the Germline Exome Best Practices Pipeline (DePristo et al., 2011) in GATK4 (McKenna et al., 2010). Briefly, we aligned reads to the reference sequences, and merged these with unaligned reads to identify duplicate read clusters based on read mapping rather than sequence identity (GATK MarkDuplicates). For each individual, we called provisional variants separately (GATK HaplotypeCaller in GVCF mode) and then called genotypes jointly for all individuals in a species (GATK JointGenotypeCaller) (Poplin et al., 2017). This produced an initial variant file containing both indels and SNPs, but only the latter were retained for genetic diversity analysis. We employed a hard filtering was to remove SNPs only supported by low base quality or depth of coverage. Before calculating statistics, we generated a reduced SNP dataset to remove all SNPs with missing data within a species using PLINK (Purcell et al., 2007). Some population genetics analysis requires the use of unlinked SNPs, so a second unlinked dataset was generated for each species by filtering SNPs exceeding a variance inflation factor (VIF) using a window (5 kb), variant count (5 ct) and linkage threshold (--indep 50 5 2).

### Population genetics statistics

We calculated heterozygosity and inbreeding coefficients on the unlinked SNP datasets for each species, using PLINK. We estimated Tajima’s D, which accounts for allelic variation, on the larger SNP dataset (linked SNPs but no missing data) using vcftools (Danecek et al., 2011). Critical values for detecting deviation from mutation-shift balance with four individuals were taken from Tajima (1989). All scripts used to conduct variant detection and population genetics statistics are freely available at github.com/lindsawi/HybSeqSNPExtraction

## RESULTS

Gene recovery was successful in all species, although gene recovery rate (measured as 25% of the targeted gene) was variable— from an average of 64 genes recovered in *Bothriochola sprinfieldii* to 321 genes in *Nerisyerinia camporum* (Figure 3C, Supplemental Table S1). We found that gene recovery was typically above 200 genes when the number of mapped reads exceeded 25,000 (Figure 3B). Target efficiency (percentage of reads mapping to target loci in HybPiper) ranged between 5% and 20%, except for *Neriyserinia camporum*, which had an average target efficiency of 43%. While much of the DNA extracted from the herbarium specimens was degraded, a third of the samples required further fragmentation before library preparation to increase number of fragments in our desired size range (~500 bp). We found no significant difference in target efficiency between fragmented and non-fragmented samples (Welsch Two Sample t-test t(42.7)= −1.07, p = 0.292).

**Figure 3:**
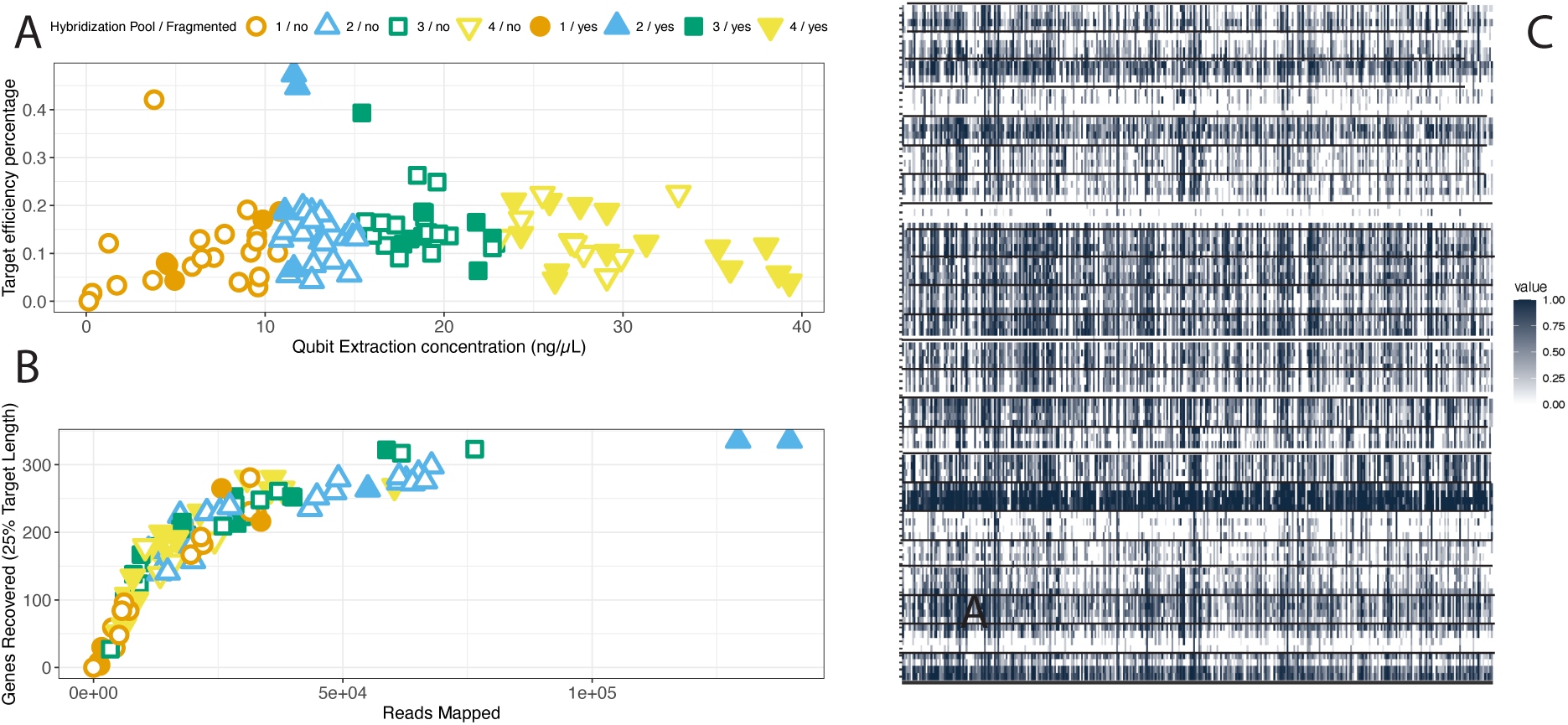
A: Relationship between sequencing library concentration (measured on a Qubit fluorometer) and target capture efficiency (percentage of reads on target). B: Relationship between gene recovery (at least 25% of the targeted gene recovered) and the number of reads on target. C: Heatmap showing the recovery of each gene (column) for each sample (row) organized by species. The shading of each cell shows the percentage of the targeted gene length recovered.

Library concentration and pooling strategy leading into hybridization reactions were among the biggest contributors to target enrichment efficiency (Fig. 3). Pool 1, which had the lowest library concentrations (0.139-10.8 ng/ul) also had the lowest target capture efficiency and had the most samples with poor gene recovery (< 100 genes, Fig. 4). In each of the other pools, there was no association between target efficiency and library concentration. While low DNA and library concentrations will not lead to optimal target efficiency, it is possible to obtain hundreds of gene sequences from these samples (Fig. 3). However, it is important that these samples have equal representation compared to higher concentration samples when pooling by equalizing molarity between sample inputs. We pooled based on concentration alone, and while the concentrations of libraries within pools 2, 3, and 4 were fairly close in value, pool 1 contained a few samples that were clearly underrepresented and resulted in poor enrichment efficiency and gene recovery (Fig. 3).

**Figure 4:**
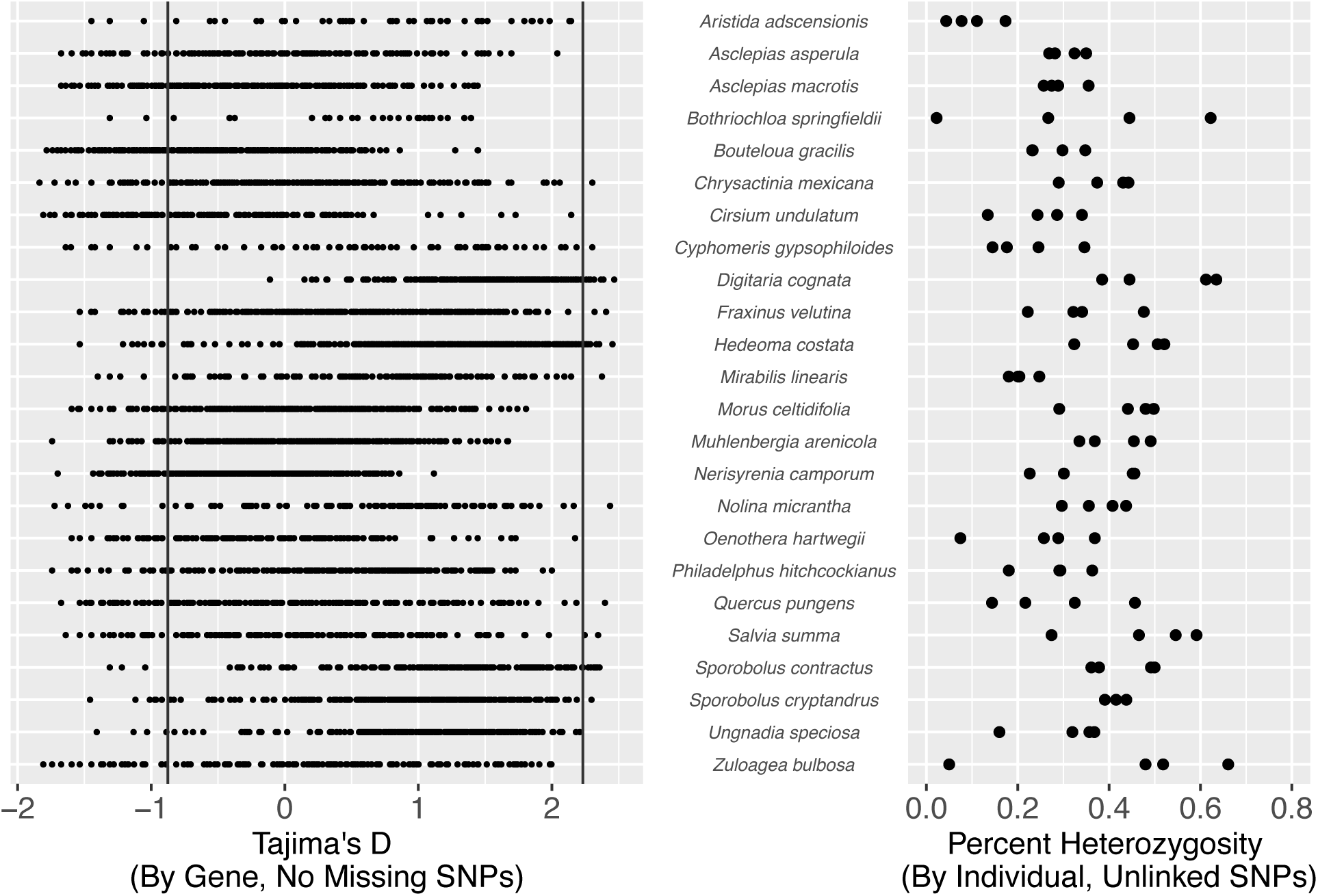
Left: Tajima’s D statistic within each species for each gene recovered, using SNPs passing hard filter and having no missing data. Vertical lines indicate 95% confidence intervals for Tajima’s D calculated for four individuals. Right: Percent heterozygosity for each individual at Angiosperms353 loci, calculated using a subset of SNPs that passed hard filtering, had no missing data, and lacked evidence of linkage disequilibrium (unlinked SNPs).

We used the longest gene recovered for each species by HybPiper to serve as a reference sequence. Notably two species, *Zuloagea bulbosa* and *Quercus pungens* contained individual supercontigs (coding sequence and flanking non-coding sequence) with lengths over 10,000 bp. We retrieved over 5000 bp for an additional fourteen species (Table 1). Total supercontig length from HybPiper was high compared to the total region targeted region of Angiosperms353 (260 kbp of coding sequences). Three species (*Nerisyrenia camporum*, *Ungnadia speciosa*, and *Zuloagea bulbosa*) had a total supercontig length over 500 kbp, most notably *N. camporum* (637216 bp). Only one species did not recover supercontigs greater than 100 kbp: *Cyphomeris gyposphiloides*.

**Table 1.**
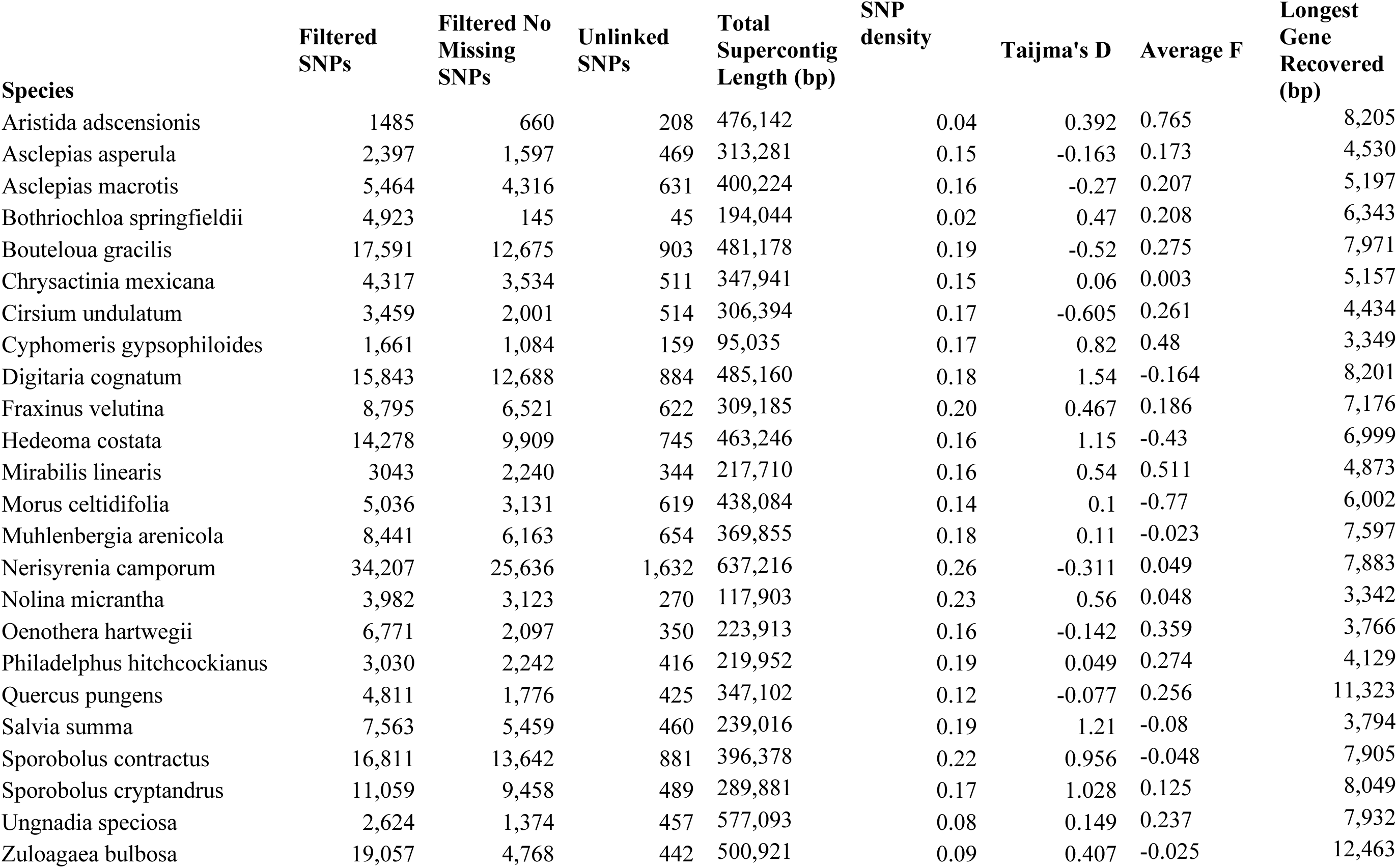
SNP and Population statistics for 24 species.

| **Species** | **Filtered SNPs** | **Filtered No Missing SNPs** | **Unlinked SNPs** | **Total Supercontig Length (bp)** | **SNP density** | **Tajima's D** | **Average F** | **Longest Gene Recovered (bp)** |
| --- | --- | --- | --- | --- | --- | --- | --- | --- |
| *Aristida adscensionis* | 1485 | 660 | 208 | 476,142 | 0.04 | 0.392 | 0.765 | 8,205 |
| *Asclepias asperula* | 2,397 | 1,597 | 469 | 313,281 | 0.15 | -0.163 | 0.173 | 4,530 |
| *Asclepias macrotis* | 5,464 | 4,316 | 631 | 400,224 | 0.16 | -0.27 | 0.207 | 5,197 |
| *Bothriochloa springfieldii* | 4,923 | 145 | 45 | 194,044 | 0.02 | 0.47 | 0.208 | 6,343 |
| *Bouteloua gracilis* | 17,591 | 12,675 | 903 | 481,178 | 0.19 | -0.52 | 0.275 | 7,971 |
| *Chrysactinia mexicana* | 4,317 | 3,534 | 511 | 347,941 | 0.15 | 0.06 | 0.003 | 5,157 |
| *Cirsium undulatum* | 3,459 | 2,001 | 514 | 306,394 | 0.17 | -0.605 | 0.261 | 4,434 |
| *Cyphomeris gypsophiloides* | 1,661 | 1,084 | 159 | 95,035 | 0.17 | 0.82 | 0.48 | 3,349 |
| *Digitaria cognatum* | 15,843 | 12,688 | 884 | 485,160 | 0.18 | 1.54 | -0.164 | 8,201 |
| *Fraxinus velutina* | 8,795 | 6,521 | 622 | 309,185 | 0.20 | 0.467 | 0.186 | 7,176 |
| *Hedeoma costata* | 14,278 | 9,909 | 745 | 463,246 | 0.16 | 1.15 | -0.43 | 6,999 |
| *Mirabilis linearis* | 3043 | 2,240 | 344 | 217,710 | 0.16 | 0.54 | 0.511 | 4,873 |
| *Morus celtidifolia* | 5,036 | 3,131 | 619 | 438,084 | 0.14 | 0.1 | -0.77 | 6,002 |
| *Muhlenbergia arenicola* | 8,441 | 6,163 | 654 | 369,855 | 0.18 | 0.11 | -0.023 | 7,597 |
| *Nerisyrenia camporum* | 34,207 | 25,636 | 1,632 | 637,216 | 0.26 | -0.311 | 0.049 | 7,883 |
| *Nolina micrantha* | 3,982 | 3,123 | 270 | 117,903 | 0.23 | 0.56 | 0.048 | 3,342 |
| *Oenothera hartwegii* | 6,771 | 2,097 | 350 | 223,913 | 0.16 | -0.142 | 0.359 | 3,766 |
| *Philadelphus hitchcockianus* | 3,030 | 2,242 | 416 | 219,952 | 0.19 | 0.049 | 0.274 | 4,129 |
| *Quercus pungens* | 4,811 | 1,776 | 425 | 347,102 | 0.12 | -0.077 | 0.256 | 11,323 |
| *Salvia summa* | 7,563 | 5,459 | 460 | 239,016 | 0.19 | 1.21 | -0.08 | 3,794 |
| *Sporobolus contractus* | 16,811 | 13,642 | 881 | 396,378 | 0.22 | 0.956 | -0.048 | 7,905 |
| *Sporobolus cryptandrus* | 11,059 | 9,458 | 489 | 289,881 | 0.17 | 1.028 | 0.125 | 8,049 |
| *Ungnadia speciosa* | 2,624 | 1,374 | 457 | 577,093 | 0.08 | 0.149 | 0.237 | 7,932 |
| *Zuloagaea bulbosa* | 19,057 | 4,768 | 442 | 500,921 | 0.09 | 0.407 | -0.025 | 12,463 |

Our data workflow (Fig. 2) resulted in a high number of variants detected within species, even for species where gene recovery was poor. Seven species initially had over 9000 variants detected from just four individuals per species, prior to filtering with PLINK. Eleven species were found to have fewer than 5000 variants, including *Aristida adscensionis* (1485 variants) and *Asclepias asperula* (2397 variants). However, unlike the poor gene recovery noted above for *Bothrichola*, the *Aristida* samples had an average of 177 genes recovered. Because many of the called variants are linked, analysis of heterozygosity within species was assessed at a subset of SNPs that contained no missing data, with one SNP chosen per LD block identified by PLINK. Average percent heterozygosity was high (above 20%) in most species (Fig. 4) and was highest in several widespread grass species including Digititaria *cognata* and *Zuloagea bulbosa*. Interestingly, heterozygosity was also high for the rare forb *Salvia summa* (a G3 vulnerable species endemic to Texas & New Mexico), and was lowest for *Aristida adscensionis*.

*Nerisyrenia camporum* contained the most variants of any of the species studied in the GUMO collection with over 34,000 SNPs recovered. After filtering, *N*. *camporum* still contained 25636 variants with 1,632variants determined to be unlinked. Notably, *Digitaria cognatum* had 12688 variants after filtering, of which 884 were unlinked, and *Sporobolus contractus* contained 13642 variants, containing 881 unlinked SNPs Additionally, SNP density was calculated to be highest for *N. camporum* (.253%), within the GUMO collection. Two grasses, *Nolina micrantha* and *Sporobolus contractus* contained the second and third highest SNP densities respectively despite low gene recovery. *Bothriochloa springfieldii* contained the lowest SNP density (023%) where only 145 filtered SNPs were detected, and only 45 were determined to be unlinked *Aristida adscensionis* contained the second lowest SNP diversity (0.43) and only contained 660 variants, of which 208 were filtered. Tajima’s D is a measurement of sequence variability that can be an indicator of deviation from mutation-drift balance (Tajima, 1989). For most of species, many Angiosperms353 loci show significantly negative Tajima’s D, which can be evidence of a recent selective sweep. With a sample of just four individuals, the critical values needed to show significant (95%) deviation from mutation-drift balance (D = 0) are –0.876 and 2.232. In our data, a much larger number of loci had significantly negative Tajima’s D compared with significantly positive (Fig. 4). At the species level, sixteen species were found to have average Tajima’s D>0. Notably, four species (*Digitaria cognata*, *Sporobolous cryptandrus, Hedeoma costata*, and *Salvia summa*) revealed an average Tajima’s D value of over 1.0. indicative of strong balancing selection or recent population contraction. We found several species with Tajima’s D<0, including *Nerisyrenia camporum*.

Inbreeding coefficients (F) calculated for each individual were averaged across individuals in each species. *Morus celtidifolia* had the lowest value (−0.77), while *Aristida adscensionis* (0.765) and Mirabilis linearis (0.511) had the highest inbreeding coefficients of the GUMO collection. Seven species were found to have negative F values (Table 1).

## DISCUSSION

### Recovery of Angiosperms353 loci

Gene recovery did not have an obvious taxonomic bias, although target capture efficiency was highest in *Nerisyrinia camporum*. This species, which is in the Brassicaceae, may have benefited from the probe design of Angiosperms353, which includes a representative from *Arabidopsis thaliana* for each gene (Johnson et al., 2018). Two of the species with the poorest gene recovery were both grasses (*Aristida adscensionis* and *Bothriotrichola springfeildii*); we believe this low recovery does not relate to poor recovery in Poaceae, as a large number of loci were recovered from the grasses *Bouteloua gracilis* and *Sporobolous contractus*. Rather, low concentration libraries in *Aristida* and *Bothritrichola* and low input DNA for several samples (Fig. 3) are the more likely explanations for low recovery. For future studies we suggest that low concentration libraries (< 5 ng/µL) should not be pooled for target capture with higher concentration libraries. Greater effort should be taken to maximize DNA input when possible, and this could include employing alternative DNA extraction techniques for recalcitrant samples (Hale et al., 2020). The poor relationship between library concentration and enrichment efficiency (for Pools 2, 3, and 4) suggests that pooling via library concentration is sufficient and calculating molarity is not necessary. Efficient recovery of more than 200 loci from each sample in HybPiper could be achieved with as few as 25,000 reads on target (Fig. 3). Given that enrichment efficiency was typically around 10-15%, it suggests 250,000 reads would be necessary to ensure high gene recovery. With newer sequencing technologies including the Illumina NovaSeq SP, which has an output of 400M reads per lane, our results suggest that massive pooling of samples (at least 1500, given flexible library indices) could be achieved.

### Genetic variability within species

Estimates of heterozygosity within species should not be taken as fully representative of the GUMO populations – with only four specimens the minimum minor allele frequency we could detect is 12.5%. However, the values can be indicative of the amount of within-species genetic diversity that can be expected from both protein coding and flanking non-coding regions of the Angiosperms353 genes. Even when using conservative filters – removing all missing data and retaining only SNPs with no evidence of linkage – hundreds of SNPs were variable within most species. We identified high heterozygosity for several species where low genetic variation might be expected, such as *Salvia summa* (a G3 vulnerable species) and other range-restricted species (e.g. *Philadelphus hitchcockianus*, *Nerisyrinia camporum*, and *Quercus pungens*).

Significantly negative values of Tajima’s D could be found for some loci in most species. As this could be evidence of selective sweeps, SNPs from these loci should be removed from analysis in future genetic studies. Species level patterns are difficult to interpret with few individuals sampled, but several species did have an average Tajima’s that was positive, potentially an indicator of population contraction or balancing selection. Notably, nearly all loci for *Digitaria cognata* showed positive Tajima’s D. Because genome-wide balancing selection seems unlikely, this result suggests recent population contraction. Although *D. cognata* is among the most widespread species in our study, GUMO is at the edge of its range, and it is not often observed at the altitudes found here (< 1800 m).

Few prior estimates of genetic diversity are available for the species selected here. The species with the most prior population-level analyses is *Bouteloua gracilis* (blue gramma): Aguado-Santacruz et al. (Aguado-Santacruz et al., 2004)and Phan (Phan, 2000) and found high variability in *B. gracilis* populations at opposite ends of its range, but these studies used RAPD markers so heterozygosity estimates could not be made. More recently, Avendano-Gonzales et al. (Avendaño-González et al., 2019)used RADSeq to characterize genetic structure across the species range and found a maximum heterozygosity of 0.32% and a Tajima’s D of –1.75. These results are in stark contrast to the values estimated here (Fig. 4), but the RADSeq study used far more individuals of *B. gracilis* (78, compared to our 4) and far more SNPs (132,000 compared to our 906). It will be interesting for future studies to directly compare RADSeq and target capture with equivalent sampling to make a better comparison between the methods. A main advantage of using Angiosperms353 loci is that studies will be more repeatable (not requiring re-discovery of loci each time new specimens are added) and will be able to incorporate longer gene haplotypes (Leitwein et al., 2020).

One study aimed at the genetic differentiation of species in *Aristida* (Thiv et al., 2019) noted little divergence of *Aristida adscensionis* from other species using traditional single-gene sequencing (*ITS*, *trnL-F*, and *rpl16*). Our results also show low genetic diversity in *A. adscensionis*, even when accounting for the low gene recovery rate. This may indicate low genetic diversity in a widespread species, and further question the universal application of the single-gene methods population level sampling. A study involving *Quercus pungens* sampling from GUMO found high heterozygosity in microsatellite markers (> 0.8), collected as part of a study aimed at identifying the origins of the federally threatened hybrid species *Quercus hinckleyi*, of which *Q. pungens* is a putative parent (Backs et al., 2016). We found slightly lower levels of heterozygosity for *Q. pungens* using the Angiosperms353 genes (Fig. 4), which may be attributed to several factors, including the difference in population parameters when comparing hypervariable markers to DNA sequences.

### Limitations

Although our results suggest a promising potential for Angiosperms353 for population genomics, we feel it is important to identify the limitations of the method. The Angiosperms353 loci were selected from a set of single-copy orthologs for all flowering plants (Leebens-Mack et al., 2019) and have an ontological bias—the genes are enriched for functions involving the chloroplast and photosynthesis (Johnson et al., 2018). As a result, their recovery may be limited in some taxa, especially those with reduced dependence on photosynthesis, including parasitic plants and mycoheterotrophs. Single-copy genes may also be more likely to be under strong purifying selection, which would limit the utility of the Angiosperms353 loci for genome wide analysis (e.g. association studies or identifying selective sweeps and regions of introgression). Depending on genomic resources, RADSeq or even whole genome sequencing may be more appropriate for genome-wide studies.

### Future Applications

Our results illustrate that Angiosperms353 can be used a cost-effective tool for estimating population parameters. Although other methods may be more appropriate depending on the goals of the study, we feel that Angiosperms353 has highest population genomics potential in two areas: conservation genomics and environmental DNA. For example, Angiosperms353 could be used to conduct population level analysis for all flowering plant species in an ecosystem without the need to develop and optimize molecular markers for each species. This application could enrich the field of conservation genomics, which is often focused on demographic parameters within threatened and endangered species, due to limited funds. However, areas like GUMO where many species are at the edge of their ranges represent special cases for biodiversity preservation. Risk of local extinction may be elevated due to low genetic diversity and distance from the rest of the species (Vucetich and Waite, 2003). Local extinction, in turn, risks the ecosystem’s health, creating cascading effects on biotic systems by removing species with critical ecosystem services (Pejchar and Mooney, 2009). Our results for *Digitaria cognata* provide a preliminary example, in which a widespread species has evidence of population contraction in an extreme environment.

Several studies have noted that comparison among related species can enhance the understanding of conservation implications drawn from population demographics (Moran et al., 1989; Barrett and Kohn, 1991; Johnson, Lang, et al., 2016). Species within the same genus could be compared with respect to differences in species ranges (e.g. in our data *Sporobolus cryptandrus* vs. *Sporobolus contractus*) or life history traits (e.g. in our data *Asclepias asperula* vs *Asclepias macrotis*). In the present study, differences among related species were limited; future investigations with deeper sampling within and among species could expand comparative analysis, to identify whether genetic diversity metrics contain phylogenetic signal. By using Angiosperms353, conservation genomics projects could ensure that sufficient data is collected for rare species while allowing for direct comparisons to the often-overlooked genetic diversity of widespread species.

Widespread application of the Angiosperms353 loci at and below the species level would enable its use as a tool for detecting plant DNA in environmental, ancient, and sedimentary samples. These metabarcoding studies frequently rely on PCR-based methods targeting very short regions of the chloroplast genome and suffer from primer universality and poor sequence differentiation among species. The results here suggest Angiosperms353 would provide ample sequence variation for identification of taxa, but would likely require the development of new bioinformatics tools to handle sequence assembly and database comparison.

## ACKNOWLEDGEMENTS

We thank L. Pokorny and E. Gardner for extensive advice in laboratory protocols. We thank the Guadalupe Mountains National Park for permission to destructively sample herbarium specimens, and M. Fokar from the Texas Tech University Center for Bioinformatics and Genomics for assistance with sequencing. MS is a Texas Tech University Honors College Undergraduate Research Scholar supported by the <u>**CH**</u> Foundation. This work was funded by contributions to MJ by the Texas Tech University College of Arts and Sciences.

## AUTHOR CONTRIBUTIONS

MS, HH, and MJ designed the study. MS and HH sampled herbarium specimens and extracted DNA. HH prepared enriched sequencing libraries. All authors conducted data analysis; MS recovered gene sequences using HybPiper, LW built the variant calling and population genetics analysis workflow with help from MJ, and HH analyzed relationships between wet lab and bioinformatics results. All authors participated in writing the manuscript, which was approved by all authors prior to submission.

## DATA AVAILABILITY

All sequencing reads are accessible on the NCBI Sequence Read Archive (SRA) under BioProject PRJNA667845. Sequences recovered for all 24 species are available on Dryad at https://doi.org/10.5061/dryad.76hdr7sv3. All herbarium specimen vouchers are described in the Appendix including TTC accession numbers. Scripts used for data analysis are available at github.com/lindsawi/HybSeqSNPExtraction

## SUPPORTING INFORMATION

**Supporting Information Table S1**. Statistics on DNA concentrations, Illumina Libraries, and HybPiper gene recovery statistics.

## Appendix 1. Voucher table for 95 herbarium specimens sampled for this study

All specimens were collected within Hudspeth and Culberson Counties in Texas, including Guadalupe Mountains National Park. All specimens are housed at the E.L. Reed Herbarium (TTC).

***Taxon***-DNA voucher information (TTC ID, *collector and number*)

***Aristida adscensionis*** L.; TTC020876 *T.L. Burgess 730*; TTC021123 *T.L. Burgess 874;* TTC020822 *T.L. Burgess 531*; TTC020821 *T.L. Burgess 638*

***Asclepias asperula (Decne.)*** Woodson; TTC021110 *T.L. Burgess 2036*; TTC020017 *T.L. Burgess 3276*; TTC020020 *T.L. Burgess 1952*; TTC020019 *T.L. Burgess 802*

***Asclepias macrotis*** Torr.; TTC020009 L.T. Green 71; TTC020011 *T.L. Burgess 1296*; TTC020010 *T.L. Burgess 2477*; TTC020012 *T.L. Burgess 2069*

***Bothriochloa springfieldii*** (Gould) Parodi; TTC020471 *T.L. Burgess 3594*; TTC020614 *T.L. Burgess 2069 3991*; TTC020890 *T.L. Burgess 827*; TTC020613 *T.L. Burgess 3995*

***Bouteloua gracilis*** (Willd. Ex Kunth) Lag. Ex Griffiths; TTC021101 *T.L. Burgess 632*; TTC021102 *T.L. Burgess 4667*; TTC021100 *T.L. Burgess 547*; TTC021099 *T.L. Burgess 838*

***Chrysactinia mexicana*** A. Gray; TTC020106 *T.L. Burgess 1494*; TTC020111 *T.L. Burgess 342*; TTC020108 *T.L. Burgess 994*; TTC020109 *T.L. Burgess 444*

***Cirsium undulatum*** (Nutt.) Spreng.; TTC021116 *T.L. Burgess 2017*; TTC021114 *T.L. Burgess 2082-1*; TTC021115 *T.L. Burgess 2031*; TTC021113 *T.L. Burgess 4545*

***Cyphomeris gypsophiloides*** (M. Martens & Galeotti) Standl.; TTC021112 *T.L. Burgess 679*; TTC020664 *T.L. Burgess 3984*; TTC020669 *T.L. Burgess 1479*; TTC020666 *T.L. Burgess 2349*

***Digitaria cognata*** (schult.) Pilg.; TTC021096 *T.L. Burgess 828*; TTC021097 *D.K. Northington 573*; TTC020956 *T.L. Burgess 655*; TTC021098 *T.L. Burgess 977*

***Fraxinus velutina*** Torr.; TTC020639 *D.K. Northington 487*; TTC020636 *T.L. Burgess 2028*; TTC020637 *T.L. Burgess 2027*; TTC020717 *T.L. Burgess 806*

***Hedeoma costata*** A. Gray; TTC020434 *T.L. Burgess 3232*; TTC020436 *T.L. Burgess 2335*; TTC020439 *D.K. Northington 436;* TTC020438 *D.K. Northington 507*

***Mirabilis linearis*** (Pursh) Heimerl; TTC021109 *T.L. Burgess 2548*; TTC021108 *T.L. Burgess 3893*; TTC020774 *T.L. Burgess 678*; TTC020771 *T.L. Burgess 4126*

***Morus celtidifolia*** Buckley; TTC020656 *T.L. Burgess 3236*; TTC020657 *T.L. Burgess 2465*; TCC021107 *T.L. Burgess 1177*; TTC020658 *T.L. Burgess 2038*

***Muhlenbergia arenicola*** Buckley; TTC021117 *T.L. Burgess 786*; TTC020921 *T.L. Burgess 830*; TTC020918 *T.L. Burgess 582*; TTC020925 *T.L. Burgess 721*

***Nerisyrenia camporum*** (A. Gray) Greene; TTC020355 *T.L. Burgess 2140*; TTC020357 *T.L. Burgess 1410*; TTC020770 *D.K. Northington 744*; TTC020359 *D.K. Northington 322*

***Nolina micrantha*** I.M. Johnst.; TTC020683 *T.L. Burgess 927*; TTC020684 *D.K. Northington 367*; TTC020804 *T.L. Burgess 4641*; TTC020681 *T.L. Burgess 2381*

***Oenothera hartwegii*** (Benth.) P.H. Raven; TTC020643 *T.L. Burgess 2488*; TTC021122 *T.L. Burgess 4486*; TTC020735 *T.L. Burgess 4405*; TTC020736 *T.L. Burgess 4457*

***Philadelphus hitchcockianus*** Hu; TTC021118 *T.L. Burgess 1142*; TTC021119 *T.L. Burgess 4243*; TTC021120 *T.L. Burgess 1955*; TTC021121 *D.K. Northington 1031*

***Quercus pungens*** Liembm.; TTC020392 *T.L. Burgess 975*; TTC021093 *T.L. Burgess 868*; TTC020391 *T.L. Burgess 1651*; TTC021094 *T.L. Burgess 4654*

***Salvia summa*** A. Nelson; TTC020496 *T.L. Burgess 3280*; TTC020502 *D.K. Northington 476*; TTC020503 *D.K. Northington 475*; TTC020497 *D.K. Northington 1032*

***Sporobolus contractus*** Hitchc.; TTC020906 *T.L. Burgess 656*; TTC020908 *T.L. Burgess 892*; TTC021105 *T.L. Burgess 537*; TTC021106 *T.L. Burgess 636*

***Sporobolus cryptandrus*** (Torr.) A. Gray; TTC021104 *T.L. Burgess 536*; TTC020900 *T.L. Burgess 722*; TTC021103 *T. L. Burgess 832*

***Ungnadia speciosa*** Endl.; TTC020570 D.K. Northington 465; TTC020569 *T.L. Burgess 1250*; TTC021095 *T.L. Burgess 1827*; TTC020567 *T.L. Burgess 4099*

***Zuloagea bulbosa*** Kunth; TTC020962 *T.L. Burgess 864*; TTC021111 *T.L. Burgess 704*; TTC020815 *T.L. Burgess 755*; TTC020461 *T.L. Burgess 3595*

